# Red blood cell classification using image processing and CNN

**DOI:** 10.1101/2020.05.16.087239

**Authors:** Mamata Anil Parab, Ninad Dileep Mehendale

## Abstract

In the medical field, the analysis of the blood sample of the patient is a critical task. Abnormalities in blood cells are accountable for various health issues. Red blood cells (RBCs) are one of the major components of blood. Classifying the RBC can allow us to diagnose different diseases. The traditional time consuming technique of visualizing RBC manually under the microscope is a tedious task and may lead to wrong interpretation because of the human error. The various health conditions can change the shape, texture, and size of normal RBCs. The proposed method has involved the use of image processing to classify the RBCs with the help of Convolution Neural Networks (CNN). The algorithm can extract the feature of each segmented cell image and classify it in various types as Microcytes, Elliptocytes, Stomatocytes, Macrocytes, Teardrop RBCs, Codocytes, Spherocytes, Sickel cell RBCs and Howell jolly RBCs. Classification is done with respect to the size, shape, and appearance of RBCs. The experiment was conducted on the blood slide collected from the hospital and RBC images were extracted from those blood slide images. The obtained results compared with reports obtained by the pathology lab and realized 98.5% accuracy. The developed system provides accurate and fast results due to which it may save the life of patients.

## 1 Introduction

Blood is the body fluid that delivers various substances such as nutritions and oxygen to cells and takes away metabolic waste from the cells. The blood accounts for 7 to 8 % of total body weight [1]. The human blood cells have three major components as platelets, white blood cells (WBCs), and red blood cells (RBCs). RBCs are the majority of counts in the blood sample, and they are accountable for providing oxygen to the various vital organ of the body, determining blood type, and also carrying away the waste product. In normal human, RBCs are flexible and appears biconcave disk-shaped with 7 to 8 μm in diameter and 2 μm in thickness. Nearly 84% of cells in the human body are RBCs. Approximately half (i.e. 40 to 45%) of the blood volume is occupied by RBCs.

According to Walkar *et al*. [2], many diseases and health conditions may be responsible for variation in shape, size, color, or appearance of RBCs.

In the case of Pyridoxine deficiency or Thalassemia or Iron deficiency, the shape of RBC shrinks compare to normal size and become almost 5 μm. This type of RBCs is termed as Microcytes. In case of severe iron deficiency, RBCs appear oval or elongated or slightly egg-shaped. Such RBCs are termed as Eliptocyes. Some liver diseases turn the shape of RBC in coffee bean-like structure and such kinds of RBCs are termed as stomatocytes. Macrocytic RBCs are the result of Vitamin B12 or Folate deficiency lead to an increase in the shape of RBC up to 9 *μ* to 14 μ. Round shaped RBCs turn into sickle-shaped due to Sickel cell anemia. Disease like Myelofibrosis and underlying marrow infiltrate change the normal RBC shape into teardrop-shaped and this kind of RBC is termed as Teardrop RBCs. Codocytes or target cells RBCs have the appearance of the shooting target with a bullseye. This target RBC is generated because of the presence of Thalassemia, Hemoglobinopathies, or liver diseases. Due to Hereditary spherocytosis and anemia, RBCs become more sphere-shaped rather than usual round, biconcave shaped. Such RBCs are termed as Spherocytes. Howell jolly RBC is the result of asplenia, in which round, dark purple to red color spot is seen in normal RBCs.

As several diseases are associated with appearance, shape, or size of RBCs, it is necessary to analyze the RBCs carefully. Traditional inspection of RBCs under the microscope is time-consuming, expensive, and needs expert knowledge. In the last few years, new technologies are working to overcome these problems.

However, this sort of only image processing technologies are facing various challenges, such as 1) RBCs can overlap each other and look like cluster which hides edges of an individual cell and make difficulties in detecting the edges. 2) The edges of RBCs may be blurry and it may due to image capturing procedure. 3) There may be less contrast between the background and foreground. This may increase the difficulties while separating out the individual RBC. 4) The artifact may present in the captured image due to light interference. To overcome all these problems we add the help of CNN to image processing. In this manuscript, we present a microscopic RBC image analysis with the Convolution neural network (CNN). CNN is a strong image classifier tool, in which image is taken as input, classify it under certain categories based on their features. In CNN, an individual unit called a neuron. Neurons are located in a series of layers. Neurons of one layer are connected to the neurons of the next layer. Each neuron or node of one layer perform mathematical calculation and pass the results to the next node. The last layer of the neural network has increased computational power due to the accumulation of experience. In the proposed technique, the colored microscopic image was converted into a grayscaled image. The Canny edge detection algorithm is used to detect edges of cells in gray-scaled images. Area-based filters are used to filter unwanted regions from the image. In this system, a single RBC image of various types is fed as an input to the deep learning network. A testing image of a single RBC cell is given as input to this trained network and checked for the presence of the disease.

## 2 Literature Review

From the last few decays, extensive work has been done in the field of blood cell segregation and analysis. This analysis is done by integrating various techniques like artificial intelligence, image processing, pattern recognition, and computer vision. Detection and evaluation of a disease can be done by studying various blood cell counts and their properties such as shape, color, size, etc. Filter-ation and ektacytometry are the standard techniques for the detection of the deformability of RBCs. Teitel *et al*. [3] worked in the same field, for quantification of RBC deformability. The technique was able to measure the mean deformability of the RBC as a population. As the technique was capable of calculating the population mean deformability it overcame the problem of analysis of individual cell deformability using a rheometer. But, at the same time, the technique was very complex and time-consuming in nature. Di *et al*. [4] used an image processing method for detection and recognition of malaria-infected red blood cells. They included morphological operators and a watershed algorithm for the processing of clustered cells. This technique provided noisy contours of RBCs. In most of the cases, due to color similarities and complex nature of blood cells, it was difficult to segment RBCs from the cytoplasm. Cai *et al*. [5] proposed a method to overcome this problem by demonstrating a statistical model-based approach. This technique was used to detect the boundary of the cell and cytoplasm. After segregating the cells from the cytoplasm, for classification of these cells based on certain features the Bayesian classifier is one of the popular techniques. Ghosh *et al*. [6] and Sinha *et al*. [7] used the Bayesian classifier technique for extracting different features of blood cells. These features included average value for color composition, area and a number of the nuclear lobes to classify the leukocytes. The system achieved 82 % accuracy for leukocyte classification. Piuri *et al*. [8] used K-Nearest neighbor to classifies the leucocytes into Basophil, Neutrophil, Lymphocyte, Monocyte, and Eosinophil. The accuracy achieved by the system was 70.6 %. Tomari *et al*. [9] proposed a system for classification of normal and abnormal RBCs by studying Artificial Neural Network (ANN) classifier. Their system achieved 82 % accuracy.

## 3 Methodology

Fig. 1 shows the process flow graph. According to figure 1 the input was a raw microscopic image received from the hospital. The received images are then pre-processed and segmented to extract single RBCs images. These single RBC images are sub-images of initial raw images. Once we obtained the subimages, the features were extracted for individual RBC images. The feature extraction technique used the following 3 features shape (Canny edge detector), size(Morphological binary operation), and color (mean RGB matrix)of each RBCs. After receiving the values of features, the normalization was carried, and finally, image classification was done. CNN is used to classify the single-segmented images into its respective classes. Now we explain each step in detail in following sections

**Figure 1:**
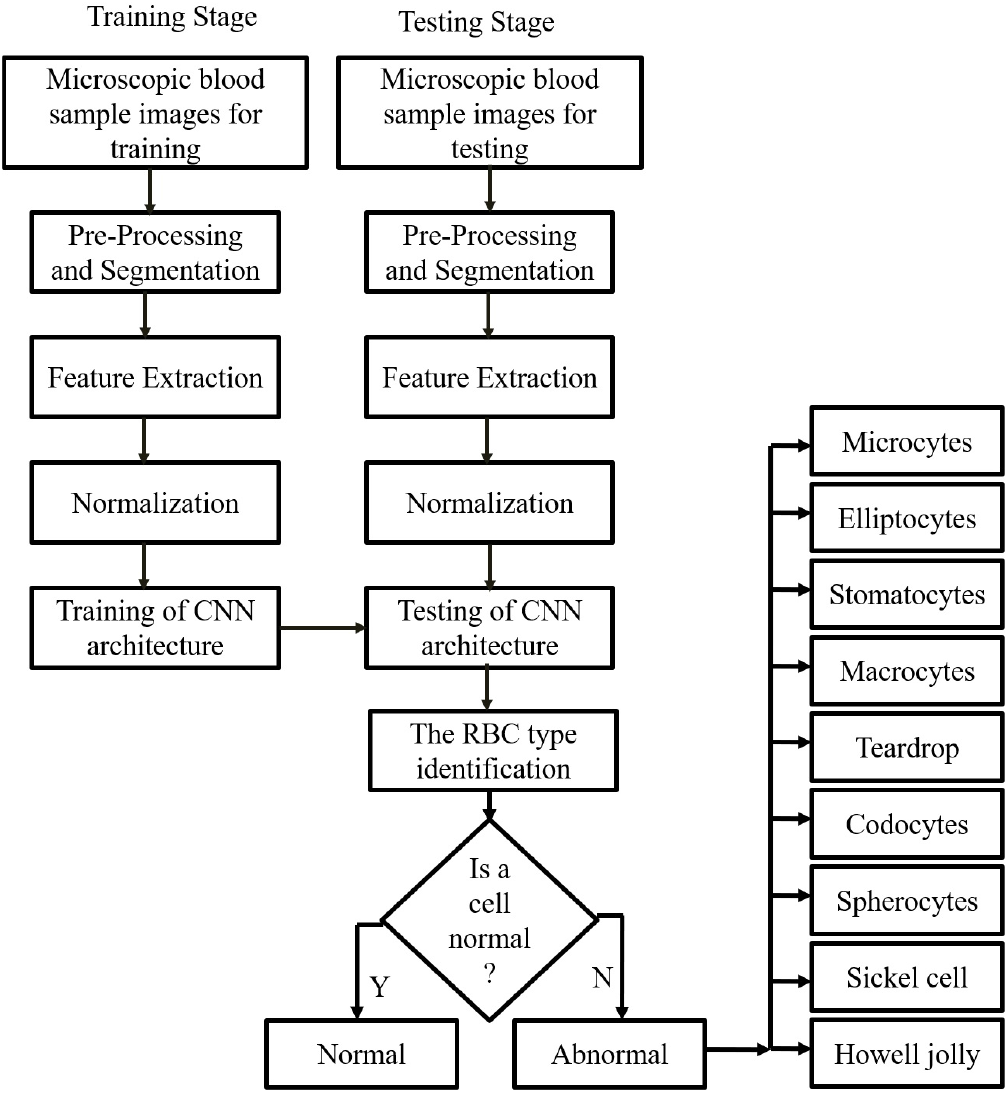
The system flow diagram.The image was acquired as a raw image from a microscope. The image pre-processing, such as, RGB to gray conversion, edge detection, ROI selection, image segmentation, normalization was performed. Once the image is pre-processed then image is normalized and features are extracted. The extracted features and original image was fed to the Convolution Neural Network (CNN) as a tensor input. CNN was trained in such a way that it was able to detect the type of RBC based on its color, shape, and texture features. The classification output was segregating RBCs into 10 classes

### 3.1 Image pre-processing

The first and essential stage of any computer vision system is image acquisition. In our case, the raw microscopic images were taken from the Olympus IX53 microscope fitted with a USB camera. The acquired blood sample images were captured from a high-resolution of 1080×1080 pixels. The acquired images were affected due to small fluctuations in light sources. To improve the dynamic range, contrast stretching was performed on the images. To specify the area taken up each cell, the image was separated into two regions as foreground and background. Rather than studying and analyzing the entire image, a small part or region of interest (ROI) around the cell was extracted and considered for further processing. Initially, for supervised learning purposes, the ROI was selected manually around the single red blood cell. Due to the extraction of ROI from the entire image, the complexity of analysis reduced also unwanted parts of the image was removed. ROI ensures that CNN can be applied to the smallest possible size to achieve easier training. Once the ROI has been selected from images, it is important to process the image to improve the quality of it. The image intensity was enhanced by histogram equalization and also edges were determined by the canny edge detector. Canny edge detection defined optimal edge finding as a set of criteria that maximize the probability of detecting true edges while minimizing the probability of false edges. A Canny edge detector uses a Gaussian convolution technique to control the degree of smoothness.

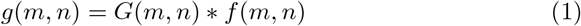

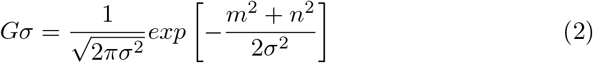

The Canny edge detection algorithm calculates the gradient magnitude and direction at each pixel. In this algorithm, the maxima and minima of the first derivative gradient are the same as the zero crossings of the second directional derivative.

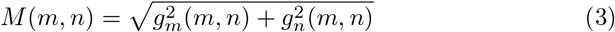

In this algorithm, the maxima crossings are taken into consideration because these pixels represent the areas of the sharpest intensity in an image. The zero-crossings are the ridge pixels that represent the set of possible edges. All other pixels are termed as non-ridge pixels and subsequently suppressed from an image. Finally, a two-threshold technique is used for performed with ridge pixels to determine the final set of pixels.

### 3.2 Feature Extraction

Feature extraction is one of the important stages of the classification of the RBC image. The shape, color, and texture are the most important features of the RBC cell image. The method used to extract shape as the feature was invariant moments, and the method applied for extracting of the texture feature was Gray-Level Co-Occurrence Matrix (GLCM). The mean, standard deviation, and skewness of pixel values are used for extracting the color feature of RBC from all three color planes.

### 3.3 CNN details

CNN is one of the simplest and effective algorithms for image classification. The CNN algorithm receives an image tensor as input and converts it into an array of pixels concatenate with features at the end. The array of such features is called a feature map. The values of the feature map depend on the resolution of the image and number of features. In the Input tensor, we used the RBC image which was normalized to 128×128×3, canny edge detected image 128×128 binary image, area, GLCM matrix, and mean R, G, B values. The CNN network is a multilayer algorithm. Convolutional, activation, ReLU, and max pooling are the most common layer of CNN. In our CNN we used 13-layer architecture (fig. 2). The first layer after tensor was a convolutional layer, also termed as moving filters (fig. 3 (c)). Convolutional layer combine feature map with convolutional filter. Due to this combination, certain features from the image get activates. After convolution, back normalization was performed. After back normalization, the Rectified linear units or ReLU (fig. 3 (a)) was used for faster training. It maps negative values to zero and maintaining positive values as it is. After 1st set of the convolutional layer (i.e. convolution, back normalization, ReLU), another 2 layers were used. Finally, max-Pooling (fig. 3 (b)) or subsampling was used. Max-pooling uses nonlinear downsampling to simplify the output. It reduces the number of the parameter that the network needs to learn. Finally fully connected CNN was used to classify the RBC image tensor into its corresponding class.

**Figure 2:**
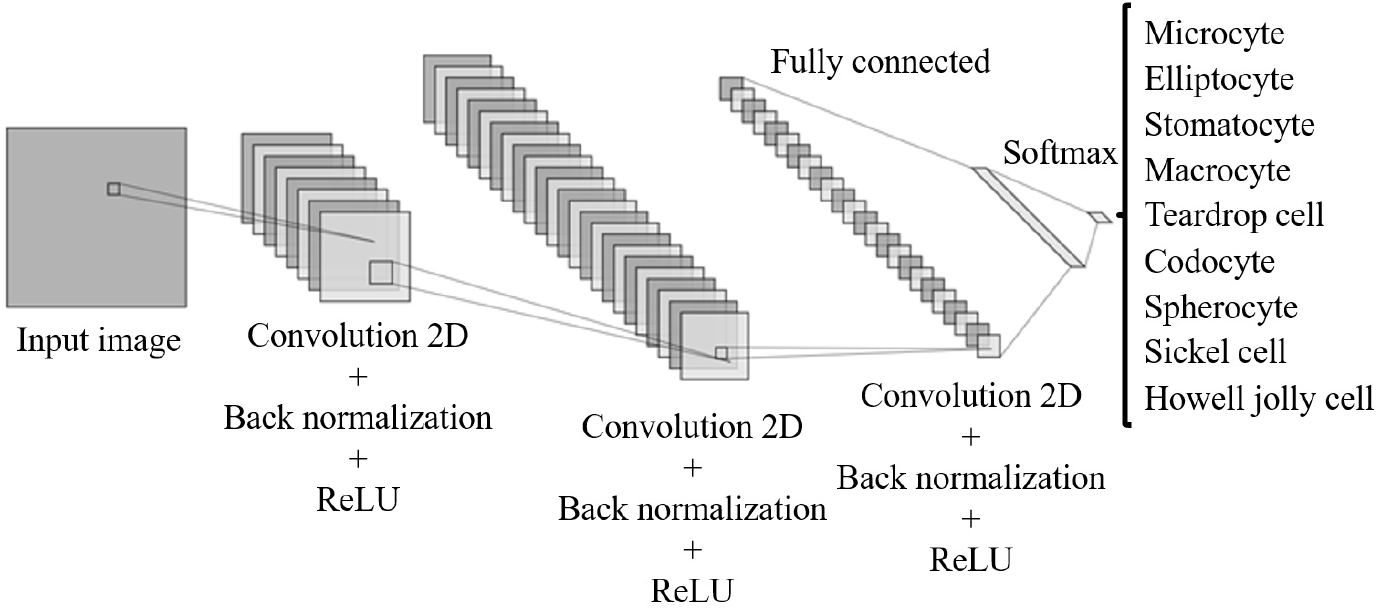
The input to CNN was tensor with an RBC image and its parts. The CNN algorithm architecture is composed of three convolutional layers with max-pooling, back normalization, and ReLU. After that, there is a fully connected network layer. A final 2-ways Softmax and classification layer provides a probability of normal and abnormal (Microcytes, Elliptocytes, Stomatocytes, Macrocytes, Teardrop RBCs, Codocytes, Spherocytes, Sickel cell RBCs and Howell jolly RBCs) per RBC image.

**Figure 3:**
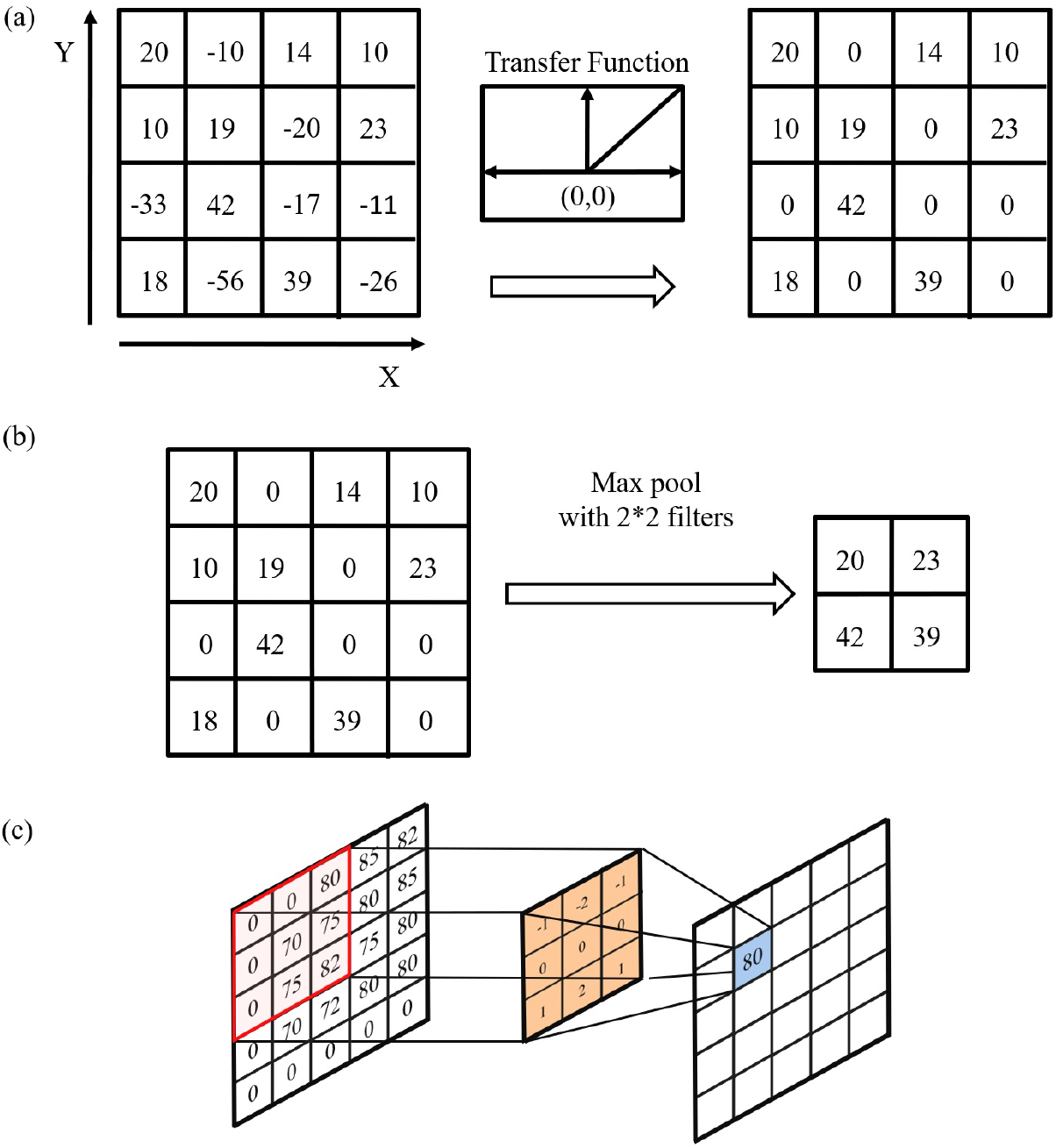
(a) Rectified linear unit (ReLU) function. It maps the negative values to zero and maintaining positive values as it is. (b) Max-pooling operation. In max pooling operation maximum value from set of 2×2 matrix is selected. (c) Filtering of image. The filter kernal of 3×3 matrix has been considered. Filter convolved with part of input (3×3 matrix) to produce new output matrix.

### 3.4 Implementation

We divided our dataset into three different parts as a testing dataset, validation dataset, and test dataset. To train CNN we marked one epoch to completed after 40 iterations. The input to the network was achieved by combining all the RBC images from the training dataset with a feature vector. After completing the training operation, for validation of the obtained model, we considered 500 images of each class from the validation dataset. The output data in such a case will be 10*500 i.e. we considered 5000 randomly selected labeled images for validation. We used randomly selected images from the testing dataset after validation is complete to cross-verify our results with pathology lab technicians results.

### 3.5 Evaluation criteria

We used the confusion matrix for the evaluation of the designed algorithm. Along with that precision, recall, F1 score, sensitivity, and selectivity were also considered.

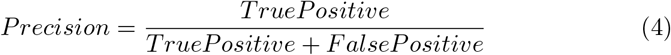

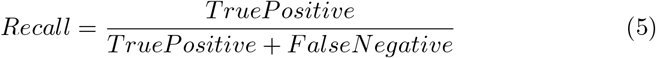

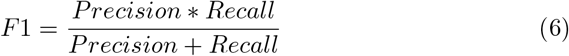

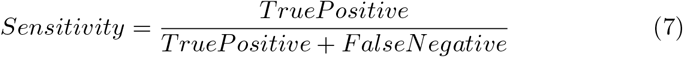

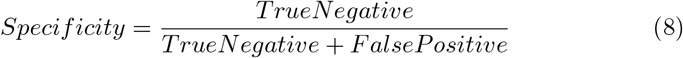

## 4 Results and discussion

Figure 4(a) Figure shows the accuracy versus total number of iteration plot. Total number of iterations are directly proportional to accuracy upto certain limit. As total number of iteration goes on increasing the accuracy of our CNN also increases. After reaching 150 iteration, accuracy saturates and become steady. After training the algorithm the accuracy of classification reached up to 98.5 % Figure 4(b) figure shows the plot of loss versus total numbers iteration. There is inverse relation between loss and total number of iteration count was observed. As the accuracy and total numbers of iteration increases the loss decreases exponentially. As shown in fig 5, the classification accuracy has reached up to overall 98.5 %. There are 10 desired classes and 10 predicted classes in the confusion matrix. The total number of RBCs in each desired class was 500 and hence we were expecting all the diagonal elements (i.e. true positive) to be 500. We got a maximum of 497/500 in teardrop cell. whereas minimum accuracy appeared in the eliptocyte class. It was expected because many of vertically oriented RBCs were looking very similar to eliptocyte, on the other hand tear drop cells have a unique shape that distinguishes it from all other classes hence chances of miss classifying was minimal. As shown in table 1 the algorithm was able to classify the image type with an overall accuracy of 98.5%. There were 500 images selected randomly from validation image folders from all 10 types of cells. As shown in the table the max True Positive (TP) count of cell type could go up to 497 out of 500. True negative could reach up to 4499 out of 4500. The maximum value of Sensitivity, Specificity, and F1 score was 99.4%, 100%, 99.4% respectively.

**Figure 4:**
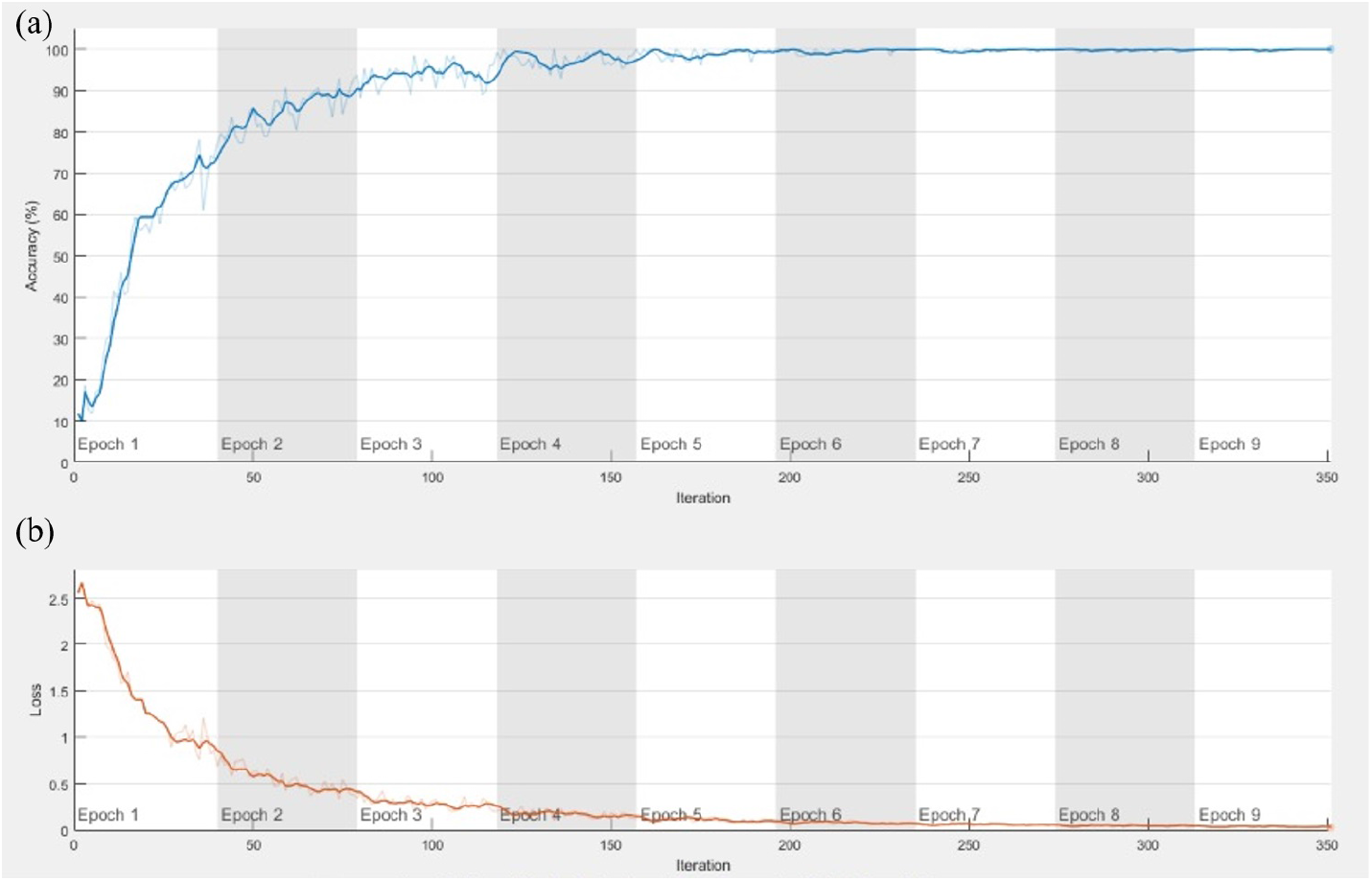
(a) The accuracy versus the number of iterations plot. The number of iteration and accuracy of CNN are directly proportional. But after certain iteration, the accuracy reaches to its saturation level. We noted that after 5 epochs, the change in accuracy was saturated. (b) shows the plot of loss versus number iterations. Loss and total number of iterations are inversely proportional. As number of iteration increases accuracy increases and loss decreases.

**Figure 5:**
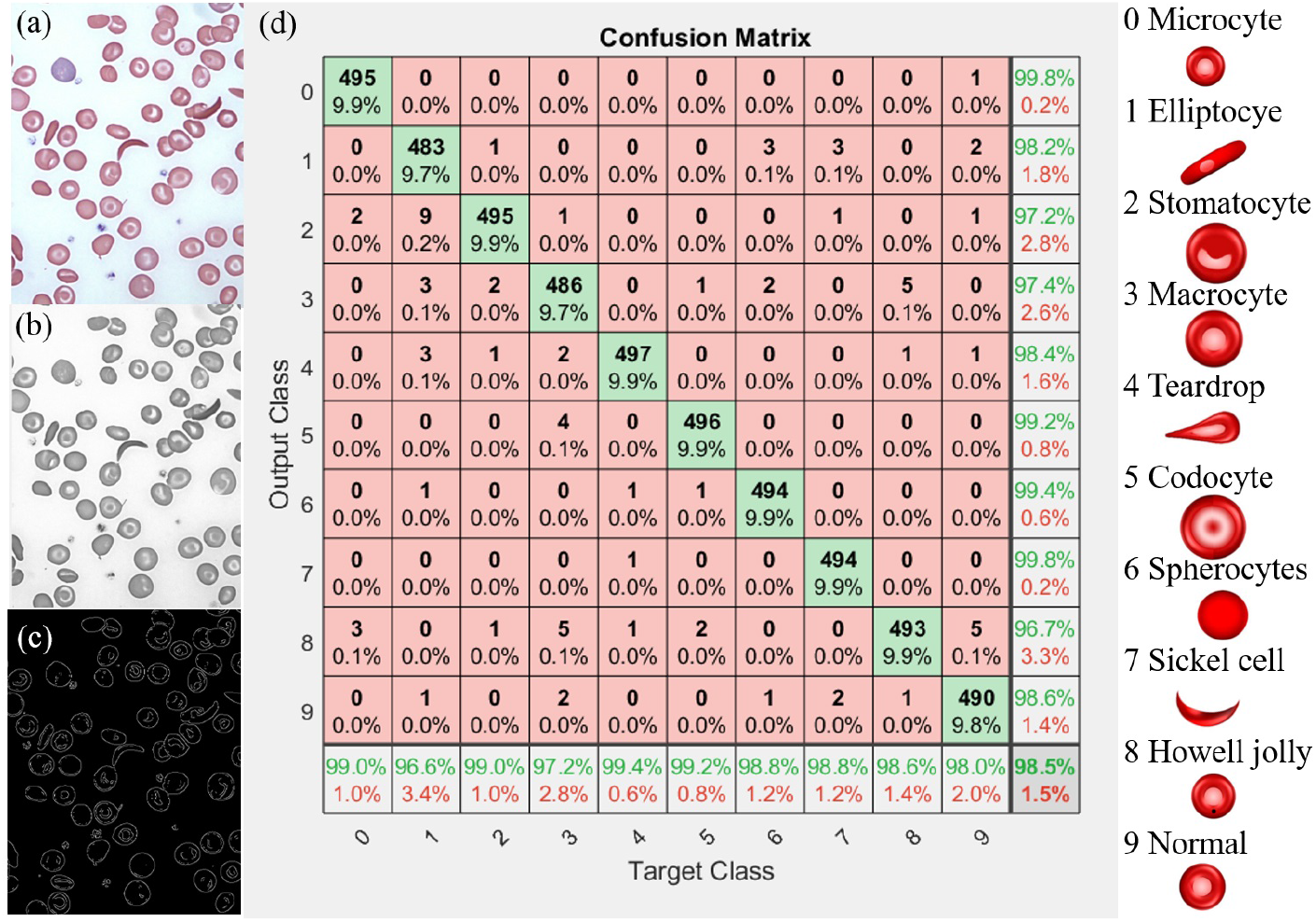
The confusion matrix obtained after classification. The input blood cell is classified into 10 types including normal RBCs and abnormal RBC of 9 types.

**Table 1:** Evaluation of CNN network

| Target Class | TP | TN | FP | FN | Precision | Accuracy | Sensitivity | Specificity | F1 Score |
| --- | --- | --- | --- | --- | --- | --- | --- | --- | --- |
| Microcyte | 495 | 4499 | 1 | 5 | 99.8% | 99.9% | 99.0% | 100.0% | 99.4% |
| Eliptocyte | 483 | 4491 | 9 | 17 | 98.2% | 99.5% | 96.6% | 99.8% | 97.4% |
| Stomatocyte | 495 | 4486 | 14 | 5 | 97.2% | 99.6% | 99.0% | 99.7% | 98.1% |
| Macrocyte | 486 | 4487 | 13 | 14 | 97.4% | 99.5% | 97.2% | 99.7% | 97.3% |
| Tear Drop | 497 | 4492 | 8 | 3 | 98.4% | 99.8% | 99.4% | 99.8% | 98.9% |
| Codocyte | 496 | 4496 | 4 | 4 | 99.2% | 99.8% | 99.2% | 99.9% | 99.2% |
| Spherocyte | 494 | 4497 | 3 | 6 | 99.4% | 99.8% | 98.8% | 99.9% | 99.1% |
| Sickle cell | 494 | 4499 | 1 | 6 | 99.8% | 99.9% | 98.8% | 100.0% | 99.3% |
| Howell Jolly | 493 | 4483 | 17 | 7 | 96.7% | 99.5% | 98.6% | 99.6% | 97.6% |
| Normal | 490 | 4493 | 7 | 10 | 98.6% | 99.7% | 98.0% | 99.8% | 98.3% |

## 5 conclusions

The RBC classification into multiclass and disease diagnostics from it is still a challenging task and it requires expert pathelogist to predict correct disease just by obervation of cells. We tried to develop an image processing and CNN based system that can detect normal and abnormal RBC with accuracy of 98% and further able to classify different types of abnormal RBCs into 9 different classes with accuracy of 98.6%. We used RBC cell images along with other features such as shape, size and texture of RBC. In the further stage of research, we are expecting to develop an deep learning method to classify RBCs from an microscopic image without any preprocessing and we expect to imaprove the accuracy of the classified results. Over all this approch can be useful not only for RBC but any muticlass segegation probleam.

## 6 acknowledgements

The authors would like to thank Dr. Madhura from Kaushalya Medical foundation hospital, Thane, and also colleagues at Ninad’s Research Lab.

## Compliance with Ethical Standards

### Involvement of human participant and animals

This article does not contain any studies with animals or Humans performed by any of the authors. And also this article contains the study on images of human blood samples. All the necessary permissions were obtained from the Institute Ethical Committee and concerned authorities.

### Information about informed consent

Informed consent was obtained from all the human participant whos blood slide images were used for image processing.

## References

[1] J.N. George, S.H. Woolf, G.E. Raskob, J. Wasser, L. Aledort, P. Ballem, V. Blanchette, J. Bussel, D. Cines, J. Kelton, et al., Idiopathic thrombocytopenic purpura: a practice guideline developed by explicit methods for the american society of hematology, Blood 88(1), 3 (1996)

[2] H.K. Walker, W.D. Hall, J.W. Hurst, Peripheral Blood Smear-Clinical Methods: The History, Physical, and Laboratory Examinations (Butterworths, 1990)

[3] P. Teitel, in Red Cell Rheology (Springer, 1978), pp. 55–70

[4] C. Di Ruberto, A. Dempster, S. Khan, B. Jarra, Analysis of infected blood cell images using morphological operators, Image and vision computing 20(2), 133 (2002)

[5] R. Cai, Q. Wu, R. Zhang, L. Fan, C. Ruan, in 2012 IEEE 11th International Conference on Signal Processing, vol. 3 (IEEE, 2012), vol. 3, pp. 1641–1644

[6] M. Ghosh, D. Das, C. Chakraborty, A.K. Ray, Automated leukocyte recognition using fuzzy divergence, Micron 41(7), 840 (2010)

[7] N. Sinha, A. Ramakrishnan, in TENCON 2003. Conference on Convergent Technologies for Asia-Pacific Region, vol. 2 (IEEE, 2003), vol. 2, pp. 547–551

[8] V. Piuri, F. Scotti, in 200f IEEE International Conference onComputational Intelligence for Measurement Systems and Applications, 2004. CIMSA. (IEEE, 2004), pp. 103–108

[9] R. Tomari, W.N.W. Zakaria, M.M.A. Jamil, F.M. Nor, N.F.N. Fuad, Computer aided system for red blood cell classification in blood smear image, Procedia Computer Science 42, 206 (2014)

